# Multi-tissue Transcriptomic and Pan-genomic Analyses Reveal Reciprocal Selective Retention Driving the Phenotypic Trade-off in Gossypium hirsutumand Gossypium barbadense

**DOI:** 10.64898/2025.12.19.693868

**Authors:** YF Xu, Zaoyang Gong, Qinglin Shen, Rongzheng Zhao, Chuankang Cheng, Wanting Su, Yibing Li, Xingrui Yang, Xin Ruan, Fengyun Li, Kai Guo, Dajun Liu, Xueying Liu, Zhonghua Teng, Fang Liu, Zhengsheng Zhang, Yanchao Xu, Dexin Liu

## Abstract

The long-standing phenotypic trade-off between Upland cotton (*Gossypium hirsutum*, high yield) and Sea Island cotton (*G. barbadense*, high quality) represents a major bottleneck in cotton breeding and industrialization. Despite rapid advances in genomics, the genomic structural variations driving the divergence of these two domestication strategies, and their subsequent impact on downstream transcriptional networks, remain unclear. This study integrates multi-tissue transcriptomic atlases spanning the full growth period with pan-genomic analyses of both Upland and Sea Island cotton, proposing a “Reciprocal Selective Retention (RSR)” strategy adopted by the two species during evolution. Transcriptomic analysis revealed that while the transcriptional chassis of the two species is highly conserved, a drastic “phase inversion” occurs during fiber development. Sea Island cotton appears to trade off for quality by specifically activating a “delayed elongation” module (at 20 DPA), whereas Upland cotton initiates a “precocious filling” program to pursue yield. Further Weighted Gene Co-expression Network Analysis (WGCNA), combined with genomic Presence/Absence Variation (PAV) analysis, identified a set of key transcription factors exhibiting a “3-vs-3” reciprocal loss pattern. Upland cotton specifically retained *CRF10*, *WIND1*, and *MYB93*, constructing a genetic foundation of “robust root system and strong regeneration” to support high yield. Conversely, Sea Island cotton specifically retained *MYB111* and *ERF105/017* to maintain long-staple characteristics and environmental buffering. Whole-genome structural variation and microsynteny analyses confirmed that these differences stem from asymmetric physical deletions (e.g., a 5 kb deletion in *MYB111*) or fragment sequence collapse (e.g., a 15.9 kb sequence divergence in *MYB93*) on homologous chromosomes. The RSR model proposed in this study not only offers a novel insight into the genetic roots of the negative correlation between yield and quality but also provides clear genomic targets for precision improvement using gene-editing technologies such as CRISPR.

## 1. Introduction

Cotton is one of the most important economic crops globally, providing the primary source of renewable natural fiber for the textile industry. Among cultivated cottons, two allotetraploid species dominate: Upland cotton (*Gossypium hirsutum*) and Sea Island cotton (*G. barbadense*) (Wen et al. 2023). Upland cotton (e.g., the standard line TM-1) accounts for over 90% of global cotton production due to its extremely high yield potential and broad environmental adaptability; however, its fiber quality (length, strength, micronaire) is relatively moderate (Avci et al. 2013). In contrast, Sea Island cotton (e.g., H7124) is renowned for its superior fiber quality, earning the reputation of “extra-long staple cotton,” but it typically suffers from lower yields and more demanding environmental requirements (Alagarsamy 2023). This phenotypic divergence between “high yield” and “high quality” reflects the distinct selection pressures the two species underwent during long-term domestication, known as “divergent domestication.” This prolonged independent evolution has led to profound divergences in their genomic structures and regulatory networks, making the combination of “high yield” and “high quality” a challenging biological trade-off to overcome in modern breeding (Liu et al. 2024).

With the rapid development of high-throughput sequencing technologies, cotton genomics research has achieved milestone breakthroughs. Since the release of the first draft genome of Upland cotton in 2015 (Li et al. 2015; Zhang et al. 2015), high-quality reference genomes for Upland cotton (TM-1) and Sea Island cotton (H7124, 3-79) have been published successively (Yuan et al. 2015; Hu et al. 2019). In particular, the recent completion of “Telomere-to-Telomere” (T2T) assemblies (Hu et al. 2025a; Hu et al. 2025b) has provided unprecedented resolution for dissecting complex allotetraploid genomes. However, despite the increasing abundance of genomic information, the precise molecular basis driving the phenotypic divergence between “high yield” and “high quality” in cotton remains incompletely resolved. Existing studies have primarily relied on Single Nucleotide Polymorphism (SNP)-based Genome-Wide Association Studies (GWAS), which often face the “missing heritability” dilemma when explaining vast phenotypic differences across species (Manolio et al. 2009). Increasing evidence suggests that, beyond minor SNPs, large-scale structural variations (such as Presence/Absence Variations, PAVs) are major drivers of adaptive evolution in polyploid crops (Pinglay et al. 2025). A current knowledge gap remains: how do these “hard” variations at the genomic level (structural variations) reshape downstream “soft” networks (tissue-specific transcriptional regulation) to establish the insurmountable “trait barriers” between Upland and Sea Island cotton (Wendel 2000)?

Cross-species comparative transcriptomics has become a powerful tool for dissecting the evolutionary mechanisms of complex traits. Yao et al. (2022) revealed the evolutionary conservation and divergence of tissue-specific expression and regulatory networks through a systematic comparison of large-scale multi-tissue transcriptomes in humans and cattle. However, this systematic multi-tissue comparative strategy has rarely been applied in the domestication research of polyploid crops. Although extensive pan-genomic studies have been conducted in the cotton field, identifying numerous structural variations associated with agronomic traits (Meng et al., 2025; Wang et al., 2022), existing comparative transcriptomic work remains largely confined to single tissues - such as fibers (Chen et al., 2012) or leaves (Cheng et al., 2025) - or specific developmental stages, lacking a systematic analysis of multi-tissue co-evolutionary patterns at the whole-plant level. Unlike diploid mammals, the domestication of allotetraploid cotton involves more complex subgenome interactions and homologous chromosome differentiation. Therefore, integrating multi-tissue spatiotemporal transcriptomics with pan-genomic structural variations is of great significance for revealing how “genomic hardware” drives the remodeling of “transcriptomic software."

In the complex transcriptional regulatory networks of plants, transcription factors serve as key bridges connecting genomic variation to phenotypic remodeling. Among them, the MYB transcription factor superfamily occupies an irreplaceable core position in cotton fiber development due to its vast membership and high functional diversity (Dubos et al., 2010; Zhang et al., 2025a). However, cotton domestication involves not only the improvement of single fiber traits but also a fundamental shift in adaptive strategies at the whole-plant level - most notably, the “broad adaptability” of Upland cotton versus the “delicate” nature of Sea Island cotton. This strongly suggests that, in addition to the MYB family, other key factors regulating cell fate determination and environmental response - such as the AP2/ERF family - may play indispensable synergistic roles in this process. AP2/ERF family members (e.g., *WIND1*, *ERF*) act not only as primary responders to hormone signals like ethylene but also as “master switches” regulating cell dedifferentiation and regeneration (Iwase et al., 2011).

Based on multi-tissue transcriptomic and pan-genomic analyses, this study reveals the molecular mechanisms driving the phenotypic divergence between Upland and Sea Island cotton. We discovered that: (1) although the global transcriptional chassis remains highly conserved, significant divergence occurs in specific tissues; (2) a distinct “phase inversion” of transcriptional programs takes place during critical stages of fiber development; (3) guided by WGCNA, we pinpointed a key regulatory module driving the “delayed elongation” trait and identified its core regulators; and (4) genomic analyses confirmed that these losses stem from asymmetric structural variations targeting genes that were originally intact in the ancestral gene pool, with pan-genomic profiling further verifying the universality of this “retention vs. loss” pattern across broad cultivated germplasm resources. Based on these findings, we propose a “Reciprocal Selective Retention (RSR)” evolutionary model in cultivated cottons, which suggests that the two species shaped their distinct adaptive strategies by asymmetrically retaining or eliminating specific functional genetic modules from the ancestral gene pool. Based on this model, we dissect the molecular logic underlying the trade-off between “high yield” and “high quality” at the level of genomic structure, providing a theoretical basis and critical genomic targets for breeding novel cotton varieties that combine both traits.

## 2. Materials and Methods

### 2.1 Plant Materials and Data Acquisition

In this study, we utilized the Telomere-to-Telomere (T2T) reference genome of Upland cotton (*G. hirsutum* TM-1) published by Yan et al. (2025), and the high-quality reference genome of Sea Island cotton (*G. barbadense* H7124) sequenced by Hu et al. (2019). The genomic data for G. barbadense were downloaded from the CottonMD database (Yang et al. 2023). Additionally, for pan-genomic evolutionary analysis, we collected chromosome-level genome assemblies of 26 Gossypium species, including wild diploids and tetraploids, from CottonGen (Yu et al. 2021) and the NCBI database (Supplementary Table S1).

### 2.2 Transcriptome Data Acquisition and Preprocessing

The gene expression data used in this study were retrieved from the CottonMD database (Yang et al. 2023). The raw high-throughput sequencing data originated from the genome sequencing project of Upland cotton (TM-1) and Sea Island cotton (H7124) by Hu et al. (2019) (BioProject: PRJNA490626). We downloaded the normalized gene expression matrix (TPM values) and selected sample data covering 21 representative tissues/stages, including roots, stems, leaves, floral organs, and ovules and fibers at various developmental stages. To enhance analysis reliability and reduce background noise, genes with extremely low expression levels (TPM < 0.1) across all samples were filtered out prior to downstream analysis.

### 2.3 Gene Expression Characterization and Comparative Transcriptomic Analysis

To evaluate global expression patterns across samples, Principal Component Analysis (PCA) was visualized using the ggplot2 package (Wickham 2016) in R (R Core Team 2025), based on a log2(TPM+1) transformed expression matrix. Additionally, t-SNE (Van der Maaten and Hinton 2008) and UMAP (McInnes et al. 2018) dimensionality reduction analyses were performed using the Rtsne and umap packages, respectively. To quantify transcriptional conservation between species, expressed genes were filtered with a threshold of TPM ≥ 0.1, and the Pearson correlation coefficient (*R*) of the number of expressed genes between corresponding tissues of TM-1 and H7124 was calculated using the “cor.test” function in R. Furthermore, to dissect divergence in expression abundance, expressed genes in each tissue were ranked by TPM values and categorized into five expression windows (Very Low, Low, Medium, High, Very High). The number of “shared genes” falling into the same expression window in both species for a given tissue was counted using custom Python scripts. Finally, to eliminate the bias of total gene number differences, we calculated the relative proportions of common genes versus species-specific genes in each tissue. For tissue specificity analysis, the Tissue Specificity Index (Tau) was calculated following the method of Yanai et al. (2005). A Tau value closer to 1 indicates stronger tissue specificity. Specific genes were defined as those with a Specificity Measure (SPM) > 0.5 (Kryuchkova-Mostacci and Robinson-Rechavi, 2017), and UpSet plots were generated using the R package UpSetR (Conway et al. 2017) to visualize the intersections of specific genes across different tissues.

### 2.4 Weighted Gene Co-expression Network Construction and Unbiased Functional Annotation

To mine key co-expression gene modules driving the delayed fiber elongation in Sea Island cotton (Avci et al. 2013), we constructed a weighted gene co-expression network using the WGCNA package in R (Langfelder and Horvath 2008). Considering computational efficiency and noise reduction, the top 6,000 genes with the highest variance in fiber development samples were selected as input data. First, a Pearson correlation matrix between genes was calculated, and a scale-free network was constructed based on a soft threshold. Subsequently, the adjacency matrix was converted into a Topological Overlap Matrix (TOM), and co-expression modules were identified using the Dynamic Tree Cut algorithm. Module expression patterns were visualized via Module Eigengenes (ME) to identify target modules specifically highly expressed in 20 DPA fibers of G. barbadense. For unbiased functional annotation of the target module, protein sequences of all genes within the module were extracted and scanned against the Pfam-A database (Mistry et al. 2021) using the hmmscan program in HMMER 3.0 software (Eddy 2011). The filtering criterion was set to an e-value < 1e-5. Based on the scan results, the types of domains contained in each gene were counted, and gene families were ranked by domain frequency to determine the most enriched categories of transcriptional regulators in the module.

### 2.5 Genome-wide Identification of Key Transcription Factor Families (MYB and AP2/ERF)

To construct a high-coverage transcription factor family atlas and mine potential unannotated loci, we performed genome-wide identification of the MYB and AP2/ERF families in the TM-1 and H7124 genomes using the Bitacora pipeline (V1.4.2) (Vizueta et al. 2020). We employed the software’s “Full Mode,” integrating genomic sequences, genome annotation files, and protein sequences for comprehensive analysis. The specific workflow was as follows: Query libraries containing Hidden Markov Models (HMM) for MYB (Pfam: PF00249) and AP2 (Pfam: PF00847) domains were constructed, with Pfam seed files downloaded from the Pfam database (Mistry et al. 2021). HMMER (Eddy 2011) was first used to scan the input reference proteomes to identify annotated family members. For Homology-based Gene Prediction, the integrated GeMoMa software (Keilwagen et al. 2016) was used to deeply mine loci potentially missed in the reference annotation. This step used homologous proteins identified in the first step as templates to predict gene structures directly from genomic DNA sequences, thereby recovering “Putative New Genes” that were missed or incompletely annotated. Subsequently, Bitacora automatically merged results from “proteome search” and “genome prediction” and removed redundancy based on coordinates. The final candidate sequences were further verified for domain integrity using the Pfam database (Mistry et al. 2021), discarding sequences with an e-value < 1e-5 or missing key domains to ensure high confidence for downstream analysis.

### 2.6 Screening and Reciprocal Validation of Species-Specific Transcription Factors

To pinpoint key candidate factors undergoing “Reciprocal Selective Retention” from vast gene families, we established a rigorous “PAV Intersection-Reciprocal Validation” screening workflow.

Initial Screening: First, genome-wide Presence/Absence Variation (PAV) lists generated by comparative genomic analysis using MUMmer4 (Marçais et al. 2018) and Minimap2 (Li 2018) were intersected with the identified MYB and AP2/ERF family members. This preliminarily screened for candidate transcription factors present in H7124 but absent in TM-1 (H7124-specific), or present in TM-1 but absent in H7124 (TM-1-specific). Reciprocal BLAST Validation: To exclude false positives caused by genome assembly errors, missing annotations, or pseudogenization, we implemented strict sequence-level validation for all candidate genes. The complete CDS sequences of candidate genes were extracted as queries and reciprocally aligned to the counterpart reference genome using BLASTN (Camacho et al. 2009) (e.g., aligning H7124-specific candidates back to the TM-1 genome, and vice versa). Retention Criteria: We retained only those genes for which no valid homologous match was detected in the counterpart genome, or where detected matches exhibited severe sequence truncation (Coverage < 60%) and high sequence divergence (Identity < 85%).

### 2.7 Genomic Structural Variation and Microsynteny Analysis

To elucidate the physical mechanisms leading to the loss of the aforementioned key transcription factors (e.g., physical deletion/NOTAL or sequence collapse/HDR), we combined whole-genome alignment with local microsynteny analysis.

Whole-Genome Structural Variation (SV) Identification: The SyRI (Synteny and Rearrangement Identifier) tool (Goel et al. 2019) was used to systematically identify syntenic regions, inversions, translocations, and local variations (including NOTAL and HDR) based on NUCmer (Marçais et al. 2018) alignment results of TM-1 and H7124 (parameters: --maxmatch -l 100 -c 1000). The Python tool plotsr (Goel and Schneeberger 2022) was used to visualize the SV status of candidate gene loci. Targeted Microsynteny Mapping: To reconstruct the evolutionary history of the six candidate loci, we selected eight representative Gossypium genomes (Supplementary Table S1) to construct microsynteny maps. Centered on the target gene (e.g., MYB111 or WIND1), 2–5 single-copy, highly conserved genes upstream and downstream were selected as physical anchors. Genomic sequences between anchors were extracted, and the presence form (intact, truncated, or completely lost) of the target gene in different species was determined using BLASTN (Camacho et al. 2009). The Python tool plotsr (Goel and Schneeberger 2022) was then used to plot microsynteny structures to illustrate the specific types of gene loss.

### 2.8 Pan-genome Homology Search and Evolutionary Trajectory Analysis

To verify the universality of the RSR pattern in Gossypium germplasm resources, we extended the analysis to 26 Gossypium pan-genomes. Using the CDS sequences of the six candidate genes as probes, homology searches were performed against all pan-genomes using the BLAST+ package (Camacho et al. 2009) (threshold: e-value < 1e-5). A custom Python script was used to extract the best hit locus in each genome and calculate its Identity and Coverage relative to the query sequence. Simultaneously, chromosomal location information (A-subgenome or D-subgenome) of the best hit was recorded to distinguish between orthologs and partially retained homeologs. The final results were visualized as bubble plots using the Python tool plotsr (Goel and Schneeberger 2022) to display the lineage-specific retention and loss trajectories of these key factors during the domestication from wild species to cultivars.

## 3. Results

### 3.1 Global Transcriptome Landscape Reveals Evolutionary Conservatism and Tissue-Specific Divergence Between Upland and Sea Island Cotton

To systematically dissect the regulatory divergence driving the phenotypic trade-off between yield and quality, we constructed a comprehensive transcriptomic atlas covering 21 tissues throughout the full growth period of Upland cotton (G. hirsutum, TM-1) and Sea Island cotton (G. barbadense, H7124).

Dimensionality reduction analysis (Fig. 1A) showed that samples clustered tightly by tissue type rather than species. Vegetative tissues (roots, stems, leaves) and floral organs of both species clustered together, indicating that despite divergent domestication, the core transcriptional chassis maintaining basic physiological functions remains highly conserved. However, beneath this global conservation, we observed significant lineage-specific differentiation in transcriptional activity. Comparison of expressed gene numbers across homologous tissues showed moderate correlation (Pearson’s R = 0.63) but highlighted unique outliers closely related to agronomic traits (Fig. 1B). Notably, TM-1 exhibited significantly more expressed genes in roots and anthers, reflecting its robust resource acquisition capability and reproductive potential. In contrast, H7124 maintained a more active transcriptome in developing fibers (especially at 20 DPA) and late-stage ovules, suggesting a more persistent regulatory program underlying its superior fiber quality. Furthermore, the distribution of the Tissue Specificity Index (Tau) revealed a shift in regulatory strategies (Fig. 1C). While both species exhibited a bimodal distribution, TM-1 showed an expansion in the number of highly tissue-specific genes (Tau > 0.5). To verify this differentiation at the gene level, we further performed pairwise comparisons of Tau indices for orthologous genes (Supplementary Fig. 3). The analysis showed that despite extremely high evolutionary conservation genome-wide (R = 0.951), confirming that functional specialization of most genes has been “canalized,” we still identified a series of “outlier genes” significantly deviating from the diagonal. These genes, which underwent drastic drift in expression breadth between species, combined with the expansion observed in Fig. 1C, strongly suggest that the evolutionary driving force for the “high yield” trait of TM-1 stems from the directional specialization and remodeling of key functional modules in specific organs (especially underground roots and reproductive sinks).

**Fig. 1.**
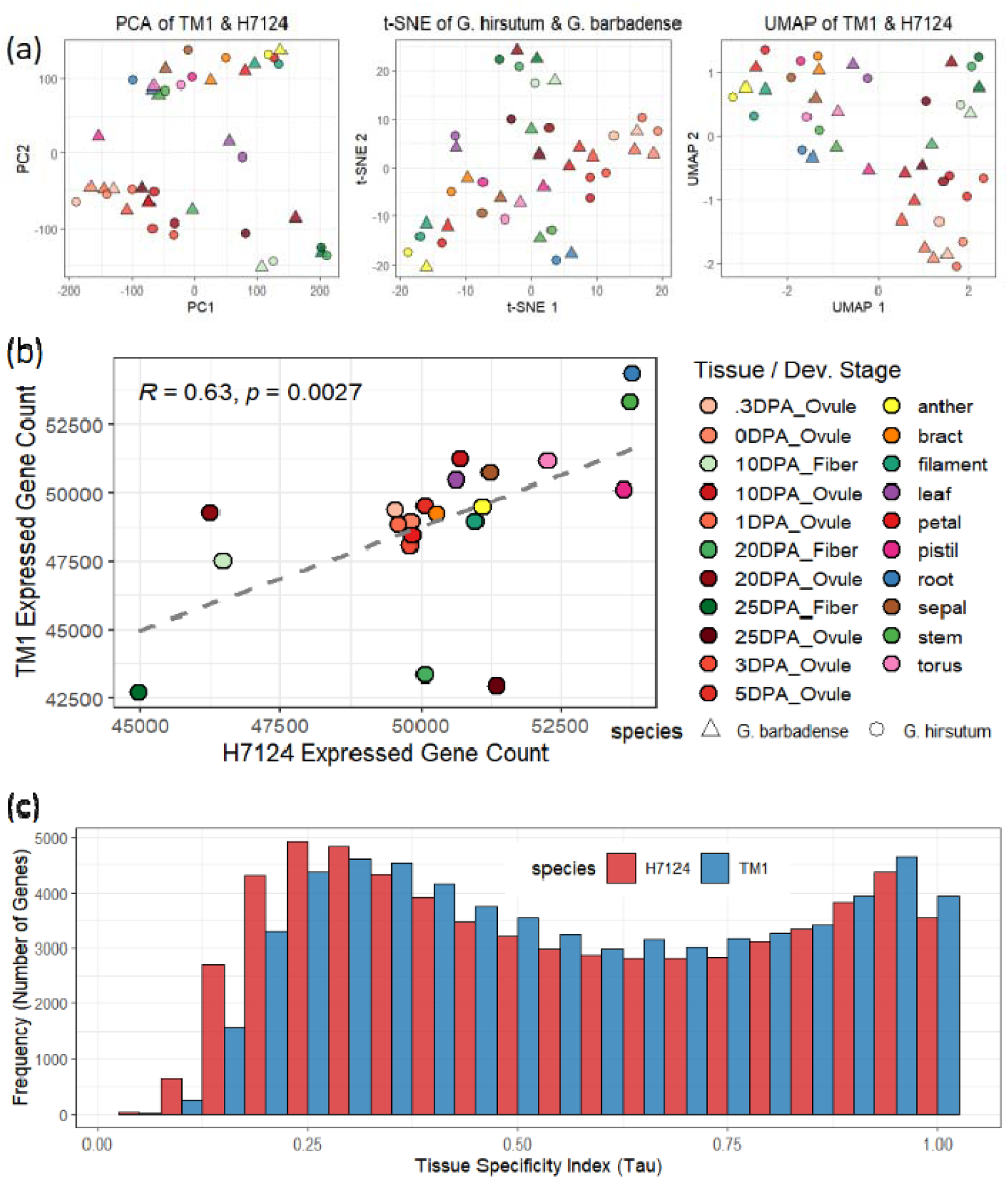
Global transcriptome landscape of Upland cotton (TM-1) and Sea Island cotton (H7124) reveals evolutionary conservation and tissue-specific divergence. (a) Dimensionality reduction analyses using Principal Component Analysis (PCA), t-SNE, and UMAP based on gene expression profiles across 21 tissues. Samples clustered mainly by tissue type (color) rather than species (shape), indicating a shared conservative Transcriptional Chassis between the two species. (b) Scatter plot comparing the number of expressed genes (TPM ≥ 0.1) in homologous tissues. Although a significant positive correlation was observed (Pearson’s = 0.0027), outliers highlight lineage-specific metabolic activity differences: TM-1 exhibits a higher number of genes in roots and anthers, whereas H7124 retains more active genes in developing fibers (e.g., 20 DPA). The dashed line represents the linear regression fit. (c) Frequency distribution of the Tissue Specificity Index (Tau). Both species display a bimodal distribution; however, TM-1 shows an expansion of highly tissue-specific genes (Tau > 0.5), while H7124 retains a higher proportion of broadly expressed genes (Tau < 0.4).

### 3.2 Asymmetric Expansion of Tissue-Specific Gene Repertoires and Phase Inversion of Fiber Development Programs

To dissect the fine-tuning mechanisms determining “yield-quality” divergence, we first systematically deconstructed the distribution patterns of tissue-specific gene repertoires (SPM > 0.5) in the two species using UpSet plots (Supplementary Figs. 2 & 3). The analysis revealed that Upland cotton and Sea Island cotton diverged in their investment directions for “functionally specialized genes” during domestication. Upland cotton (TM-1) exhibited a significant “underground-priority” strategy (Supplementary Fig. 3). Its root-specific gene number underwent an explosive expansion, constituting the most significant specialized module among all tissues, suggesting the evolution of a complex root network to support high yield potential. However, as an evolutionary trade-off, its number of specific genes significantly contracted during the middle to late stages of fiber development (10–25 DPA). In contrast, although Sea Island cotton (H7124) had fewer root genes, it maintained sustained high investment in reproductive sink organs (Supplementary Fig. 2); particularly in 20 DPA Fiber (elongation-thickening transition period) and 25 DPA Fiber, H7124 still retained an extremely vast specific gene repertoire, providing the genetic basis for the fine shaping of “extra-long staple” quality. This asymmetric distribution of gene repertoires—"TM-1 strong roots” vs. “H7124 strong fibers"—further manifested as a drastic “phase inversion” in the spatiotemporal dynamics of the transcriptome (Fig. 2):

**Fig. 2.**
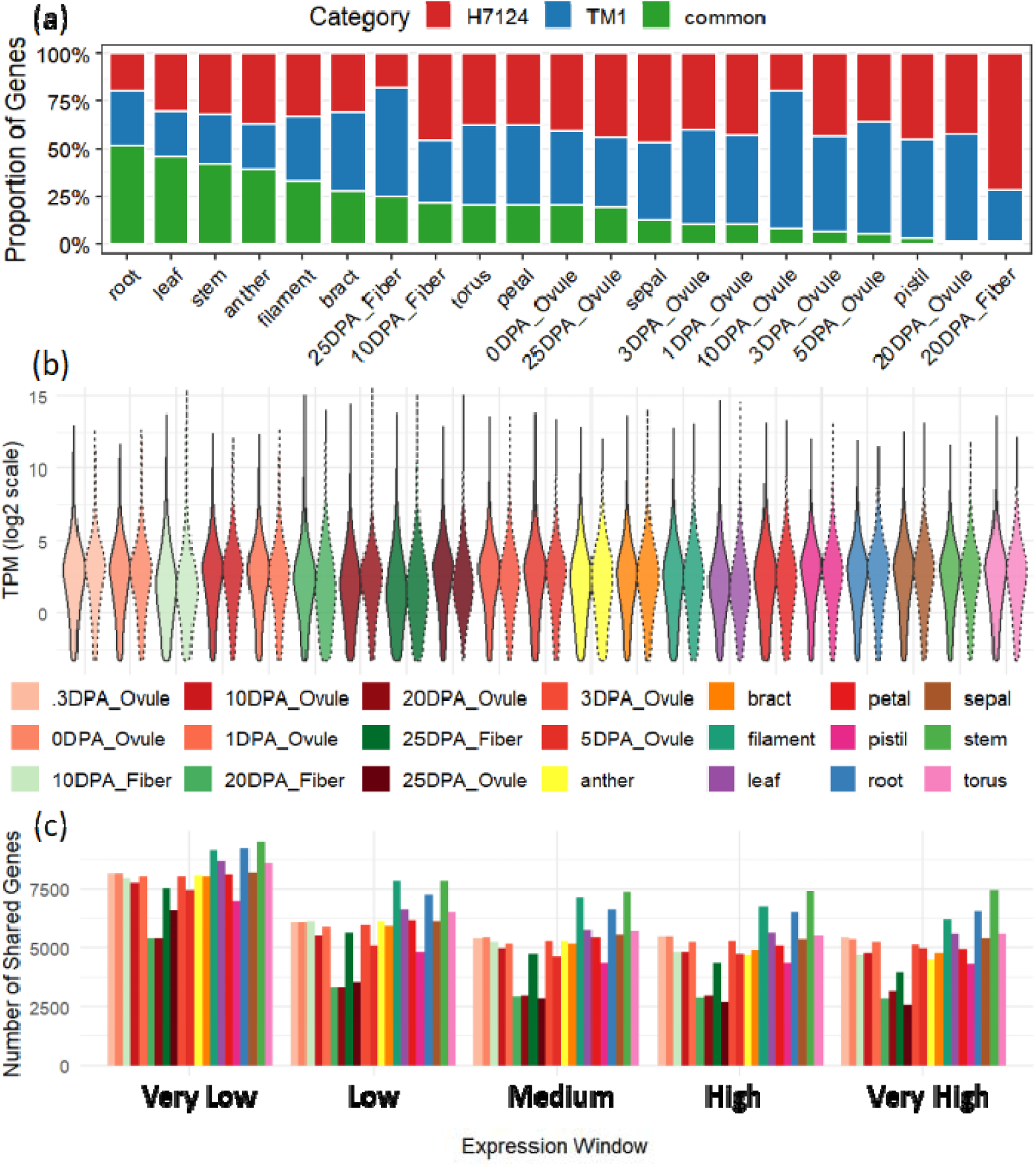
Conservation and spatiotemporal divergence in tissue-specific gene expression between Upland and Sea Island cotton. (a) Stacked bar chart showing the relative proportions of “Common” and “Species-Specific” genes (TPM ≥ 0.1) in each tissue. Green represents expressed genes common to both species, red represents H7124-specific expressed genes, and blue represents TM-1-specific expressed genes. Note the significant proportion inversion (phase inversion) occurring during fiber development (20 DPA to 25 DPA). (b) Violin plots of gene expression abundance (TPM) for H7124 (solid lines/solid fill) and TM-1 (dashed lines/hollow) across tissues. Solid lines represent H7124, and dashed lines represent TM-1. (c)Statistics of shared gene numbers within different expression abundance windows (from Very Low to Very High). The charts reveal high conservation in the transcriptomes of vegetative organs, contrasting with the drastic transcriptional remodeling in reproductive organs (particularly fibers and ovules).

First, the overall distribution of gene expression (Fig. 2b) and stratified sharing analysis (Fig. 2c) reconfirmed the evolutionary conservation of vegetative growth. In vegetative organs such as roots, stems, and leaves, the two species not only shared highly consistent overall distribution shapes of gene expression abundance but also retained a high proportion of common genes in all expression windows (especially medium-to-high expression intervals) (green bars, approx. 30%-50%, Fig. 2a), indicating that the transcriptional network maintaining basic biomass accumulation in cotton plants is stable across species.

However, entering the critical stages of fiber and ovule development, the transcriptional program underwent fundamental remodeling, presenting a significant “species-biased phase inversion” (Fig. 2a): H7124-dominated “Delayed Elongation” Phase (20 DPA): At the critical node of transition from fiber elongation to secondary wall thickening, Sea Island cotton-specific expressed genes (red bars) occupied absolute dominance (approx. 70%), while common genes and TM-1 specific genes were compressed to extremely low proportions. Combined with the wider high-expression interval of H7124 in this period in Fig. 2b, this indicates that Sea Island cotton extends the developmental window of fiber elongation by specifically activating a vast transcriptional network (i.e., the gene repertoire observed in Supplementary Fig. 2), thereby achieving the high-quality phenotype. TM-1-dominated “Precocious Filling” Phase (25 DPA): Subsequently, during the high-speed secondary wall synthesis period, the trend reversed. The proportion of Upland cotton-specific genes (blue bars) significantly rebounded and took dominance. This suggests that Upland cotton initiates the secondary wall deposition program centered on biomass accumulation earlier, reflecting its breeding selection strategy for “early maturity and high yield (high lint percentage)". In summary, although the basic transcriptional chassis of cotton is conservative, Upland and Sea Island cotton likely achieved spatiotemporal separation of developmental programs by recruiting distinct specific gene modules: H7124 chose a delay strategy of “trading time for quality,” while TM-1 chose a precocious strategy of “efficiency first". This phase inversion at the transcriptional level (Fig. 2 evidence), caused by the asymmetric expansion of gene repertoires (UpSet evidence), provides systematic molecular insights for understanding the evolutionary trade-off between “high yield” and “high quality".

### 3.3 WGCNA Reveals a “Delayed Elongation” Regulatory Module Centered on MYB-AP2

To deconstruct the “fiber development phase inversion” phenomenon observed in Fig. 2 and mine the core drivers maintaining high transcriptional activity in Sea Island cotton (H7124) at 20 DPA, we constructed a gene co-expression network using WGCNA. The analysis pinpointed Module 6 as the key target module: the expression profile of this module highly matches the “delayed elongation” phenotype of Sea Island cotton (Fig. 3a), maintaining extremely high activity in H7124 fibers at 10-20 DPA, while rapidly declining in Upland cotton (TM-1) during the same period. This significant species-specific expression pattern may imply that Module 6 is the transcriptional basis for Sea Island cotton fibers breaking through the conventional elongation time limit.

**Fig. 3.**
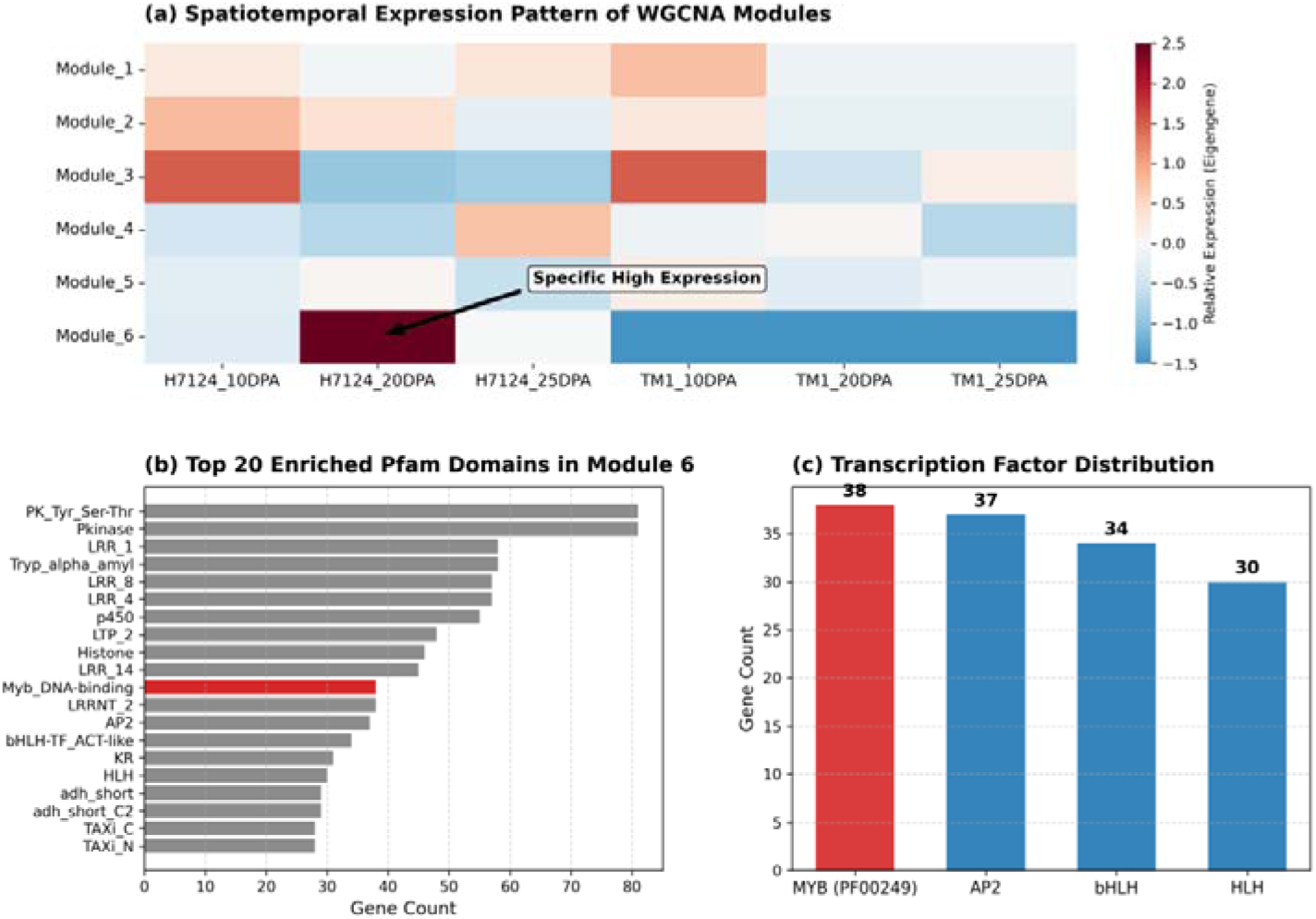
Weighted Gene Co-expression Network Analysis (WGCNA) identifies key modules driving delayed fiber elongation in Sea Island cotton. (a) Spatiotemporal expression heatmap of WGCNA Module Eigengenes. Colors represent relative expression levels (Red: High; Blue: Low). The arrow points to Module 6, which exhibits specific high transcriptional activity in H7124 20 DPA fibers, presenting a “delayed expression” characteristic. (b) Top 20 enriched Pfam domains in Module 6. This module is rich in domains involved in signal transduction (e.g., Pkinase, LRR) and transcriptional regulation, indicating that the cells are in an active state of metabolism and signal response. (c) Distribution of member counts for major transcription factor families in Module 6. The MYB and AP2/ERF families show the highest enrichment, implying their core regulatory roles in this module.

Statistics on the number of gene families contained in this module showed (Fig. 3b) that it is enriched with a large number of Protein Kinase (Pkinase) and Leucine-Rich Repeat (LRR) domains, indicating that cells are receiving continuous growth signals to maintain an active metabolic state. More critically, at the transcriptional regulation level (Fig. 3c), the MYB and AP2/ERF families showed the highest recruitment abundance. Among them, the MYB family (38 members) ranked first as recognized fiber development regulators, followed closely by the AP2/ERF family (37 members), which typically acts as ethylene response factors regulating cell elongation and stress adaptation. Considering the classic functions of the MYB family in secondary wall and fiber elongation, and the key role of AP2/ERF in coordinating hormone signals and environmental adaptation, the high co-enrichment of these two families in Module 6 implies that the Sea Island cotton-specific fiber elongation network may be orchestrated synergistically by these two types of key factors. To verify this hypothesis and trace the genetic roots leading to the expression divergence of Module 6 between species, we subsequently performed systematic identification of MYB and AP2 families genome-wide. Through intersection screening of gene family members with whole-genome PAV (Presence/Absence Variation) lists, supplemented by strict reciprocal BLAST validation (Supplementary Fig. 4 & 5), we finally locked down 6 key transcription factors that underwent “reciprocal loss” between the two cotton species from the genome. To parse the physical mechanisms of these loss events, we further performed microscopic synteny and structural variation analyses on them.

### 3.4 Reciprocal Genomic Structural Variation and Physical Loss Mechanisms of Key Transcription Factors

To dissect the physical basis of expression divergence for the key transcription factors identified by WGCNA, PAV, and reciprocal BLAST, we employed a “Reciprocal Whole-genome Alignment” strategy. First, SyRI analysis constructed a macroscopic structural variation map using each genome as a reference (Supplementary Figs. 6 & 7). Chromosome-scale visualization clearly displayed a highly collinear background (gray links) between TM-1 and H7124 at the whole-genome level (A01-A13, D01-D13), confirming the high quality and overall stability of the genome assemblies for both species. Based on this robust macroscopic framework, we precisely extracted local alignment information for the six candidate gene loci from the whole-genome alignment results and plotted high-resolution microscopic structural variation maps (Fig. 4). This progressive “zoom-in” analysis from “whole-chromosome background” to “specific loci” revealed two lineage-specific loss mechanisms:

**Fig. 4.**
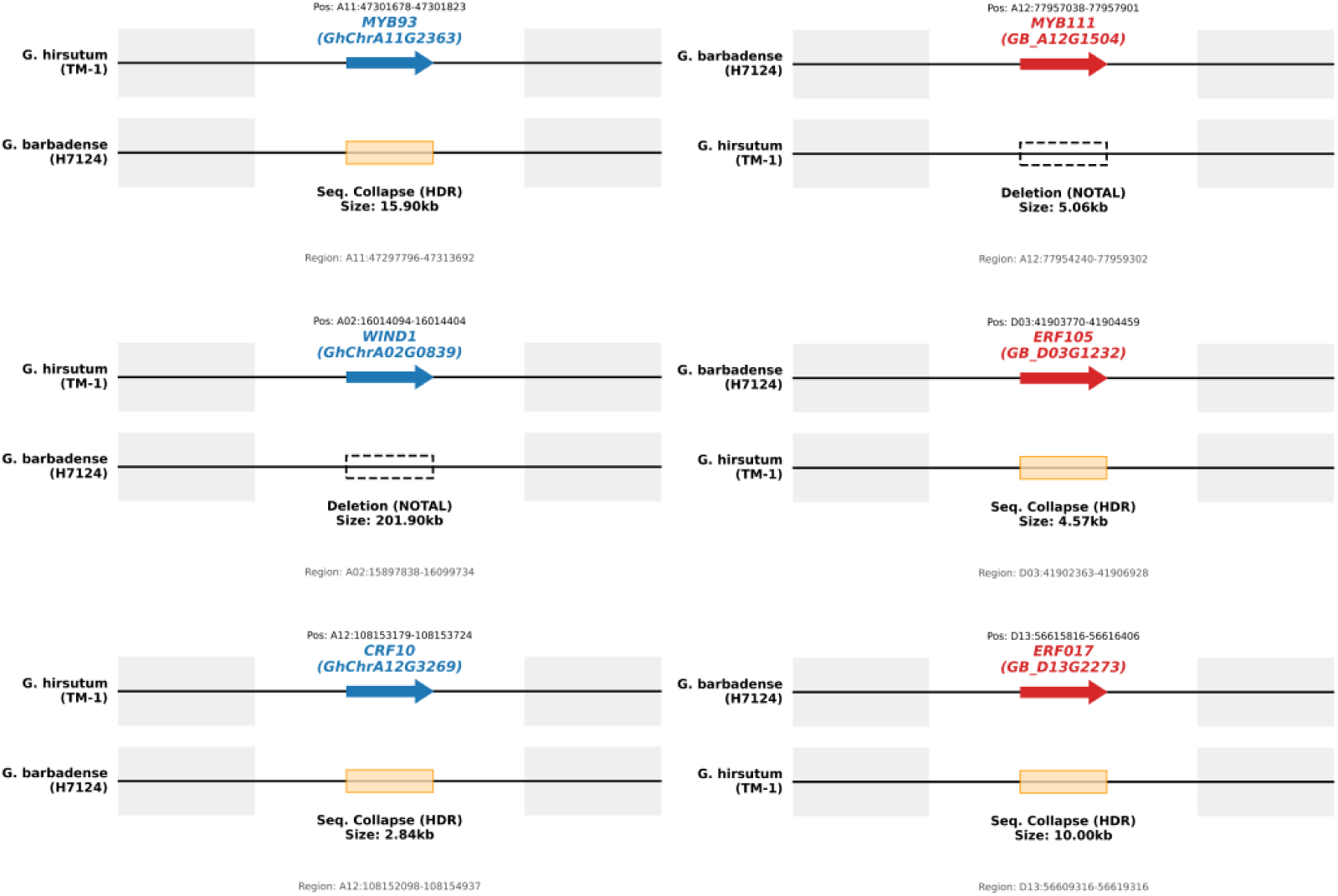
Reciprocal genomic structural variation mechanisms of key transcription factors (MYB and AP2) between Upland and Sea Island cotton. Detailed Structural Variation (SV) maps of six candidate loci based on SyRI analysis reveal two distinct lineage-specific loss mechanisms: Physical Deletion (NOTAL) and Sequence Collapse (HDR). (a) Left panel: Upland cotton (TM-1) specific retained genes (lost in H7124). Blue arrows represent intact genes in TM-1 (*MYB93, WIND1, CRF10*). Corresponding regions in the H7124 genome are identified as large-fragment deletions (e.g., a ∼201.90 kb NOTAL deletion at the WIND1 locus) or Highly Diverged Regions (HDR), resulting in the loss of functional copies. (b) Right panel: Sea Island cotton (H7124) specific retained genes (lost in TM-1). Red arrows represent intact genes in H7124 (*MYB111, ERF105, ERF017*). In TM-1, the region corresponding to *MYB111* underwent a precise ∼5.06 kb physical deletion, while *ERF105/ERF017* are located in sequence collapse regions. This genomic “Double Dissociation” constitutes the physical basis of the RSR mechanism.

First, regarding *MYB111* (A12), which is specifically retained in Sea Island cotton and serves as the core factor driving “delayed elongation” as mentioned earlier, SyRI analysis revealed the root cause of its silencing in Upland cotton: a physical deletion (NOTAL) of 5.06 kb occurred at the corresponding chromosomal position in TM-1 (Fig. 4b). This “clean” genomic excision directly removed the coding region of *MYB111*, genetically terminating the potential for further fiber elongation in Upland cotton, thereby shifting the strategy toward early maturity and high yield. In a mirror contrast, *WIND1* (A02), specifically retained in Upland cotton and a potential regulator of cell regeneration and root plasticity, underwent a catastrophic loss event in Sea Island cotton. The corresponding region in H7124 suffered a massive deletion of 201.90 kb (Fig. 4a), resulting in the complete removal of the gene and its surrounding regulatory regions. This may explain the defects in root adaptability (i.e., the “delicate” nature) of Sea Island cotton. In addition to physical deletion, Sequence Collapse (HDR) represents another major loss mechanism. For instance, the Upland-specific *MYB93* (A11) and *CRF10* (A12) both fell into Highly Diverged Regions (HDR) at their corresponding loci in Sea Island cotton, indicating that the original gene sequences underwent drastic rearrangement or degeneration, leading to loss of function.

In summary, Fig. 4 implies that this “double dissociation” phenomenon between the two cultivated species is not a sequencing error but likely a genuine genomic event. Upland cotton “discarded” quality genes like *MYB111* in exchange for early maturity, while Sea Island cotton “discarded” adaptability genes like *WIND1* in exchange for specialized development. This asymmetric structural variation on homologous chromosomes constitutes the solid genetic basis for the “yield-quality” phenotypic trade-off between the two.

### 3.5 Pan-genome Microsynteny Reveals Lineage-Specific Loss Trajectories of Key Factors

To further confirm that the structural variations identified in Fig. 4 were driven by directional selection during domestication rather than random events, we expanded our scope to the evolutionary history of the Gossypium genus. By incorporating the genomes of diploid ancestors (*G. raimondii*, *G. arboreum*) and wild tetraploids (*G. tomentosum*, *G. mustelinum*), we constructed high-resolution microsynteny maps for the six key loci (Fig. 5).

**Fig. 5.**
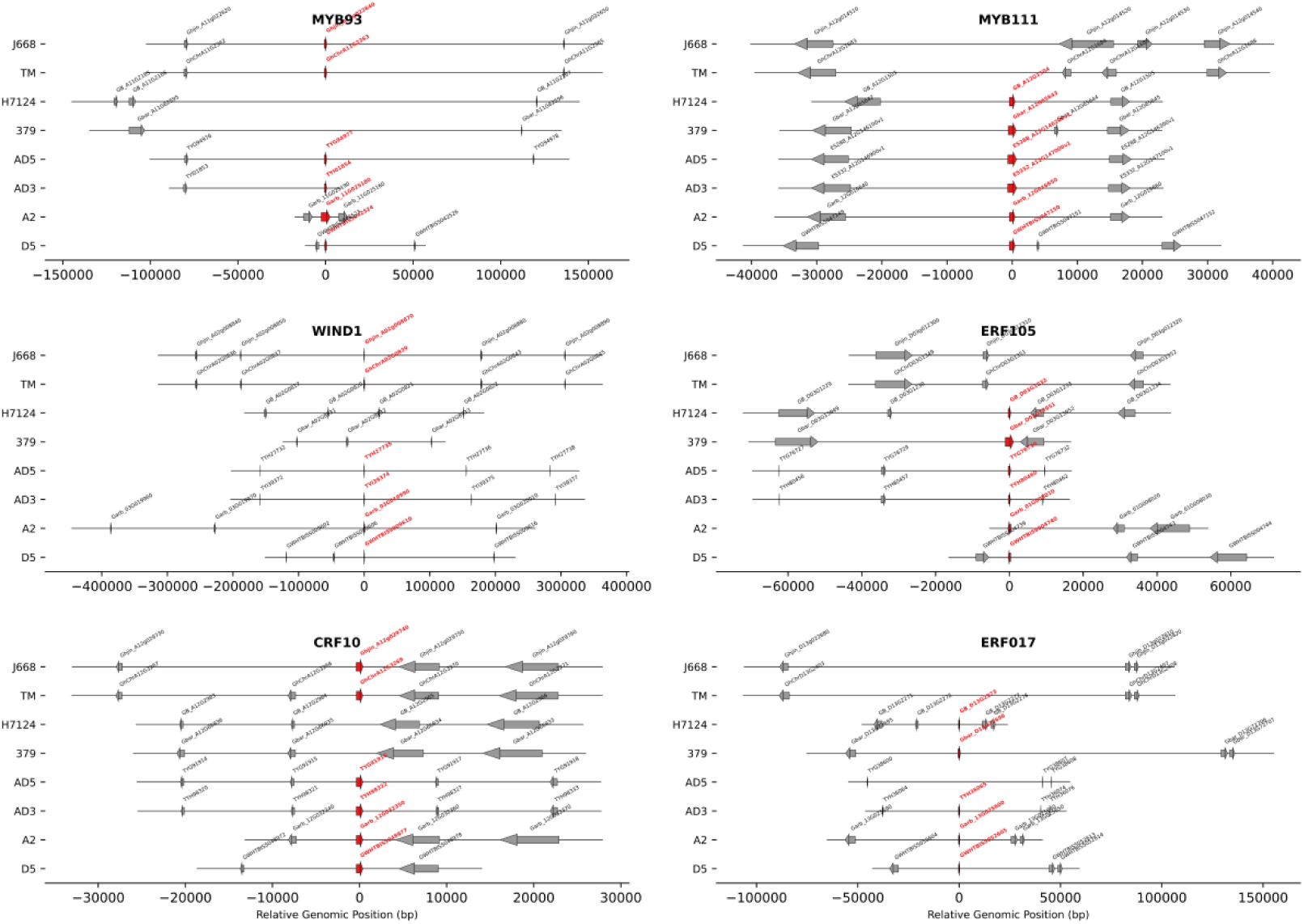
Lineage-specific loss and microsynteny maps of six key transcription factors in allotetraploid cotton. High-resolution microsynteny maps constructed using eight representative Gossypium genomes (covering diploid ancestors, wild tetraploids, and cultivars) reveal a “3-vs-3” reciprocal loss evolutionary trajectory. Group 1 (G. hirsutum-specific loss): Includes *MYB111*, *ERF017*, and *ERF105*. These genes are structurally intact and syntenically conserved (red arrows) in diploid ancestors (D5, A2) and wild tetraploids (AD3, AD5), but underwent physical deletion in the Upland cotton lineage (TM-1, J668), leaving only flanking anchor genes (gray arrows). Group 2 (G. barbadense-specific loss): Includes *MYB93*, *WIND1*, and *CRF10*. These genes are highly conserved in Upland cotton but specifically underwent collapse or large-fragment deletion (e.g., the genomic gap at the *WIND1* locus) in the Sea Island cotton lineage (H7124, 3-79). This analysis, referenced against wild species, confirms that the phenotypic differentiation of modern cultivars stems from differential purging of the ancient ancestral gene pool.

The analysis clearly revealed an evolutionary pattern of “ancestral omnipotence vs. descendant specialization,” strongly supporting the RSR hypothesis. “Upland-style Elimination” of Quality Genes: Focusing on *MYB111* (determining delayed fiber elongation) and *ERF105/ERF017* (environmental buffering factors) (Fig. 5 right panel), we found that these genes maintained intact gene structures and collinear relationships in diploid ancestors and wild tetraploids. This implies that “high quality/long staple” was originally an ancient trait potential of the Gossypium genus. However, on the divergence branch of Upland cotton (TM-1, J668), these loci underwent precise physical excision. This suggests that Upland cotton may have “actively” discarded these high-energy-consuming quality regulatory genes during domestication to achieve early maturity and high yield.

"Sea Island-style Elimination” of Adaptive Genes: Conversely, at the *MYB93* (root architecture) and *WIND1* (regeneration capacity) loci (Fig. 5 left panel), the Sea Island lineage exhibited specific loss. In particular, while WIND1 existed intact in Upland cotton and all wild relatives, a significant genomic gap appeared in H7124 and 3-79. This “functional deficiency” specifically occurring in the Sea Island evolutionary chain is likely the genetic root of its poor root adaptability and environmental sensitivity (i.e., being “delicate"). In summary, microsynteny evidence indicates that the phenotypic divergence between Upland and Sea Island cotton did not stem from the evolution of new genes, but rather from their inheritance of a complete gene pool from a common ancestor, followed by divergent “genomic subtraction” in opposite directions. This complementary trajectory of gene loss ultimately solidified their distinct agronomic traits.

### 3.6. Pan-genomic Evolutionary Footprints Confirm the Universality of Reciprocal Selective Retention (RSR)

To verify whether the RSR pattern revealed in Fig. 4 and Fig. 5 is universal across cotton germplasm resources, we extended the analysis to 26 Gossypium pan-genomes, covering diploid ancestors, wild tetraploids, and modern cultivars. Using the six key genes as probes, we constructed high-resolution phylogenomic footprints (Fig. 6). Pan-genomic alignment clearly categorized these six genes into two distinct evolutionary trajectories, revealing that the “3-vs-3” reciprocal loss pattern is likely a fixed genetic divergence at the species level between Upland and Sea Island cotton.

**Fig. 6.**
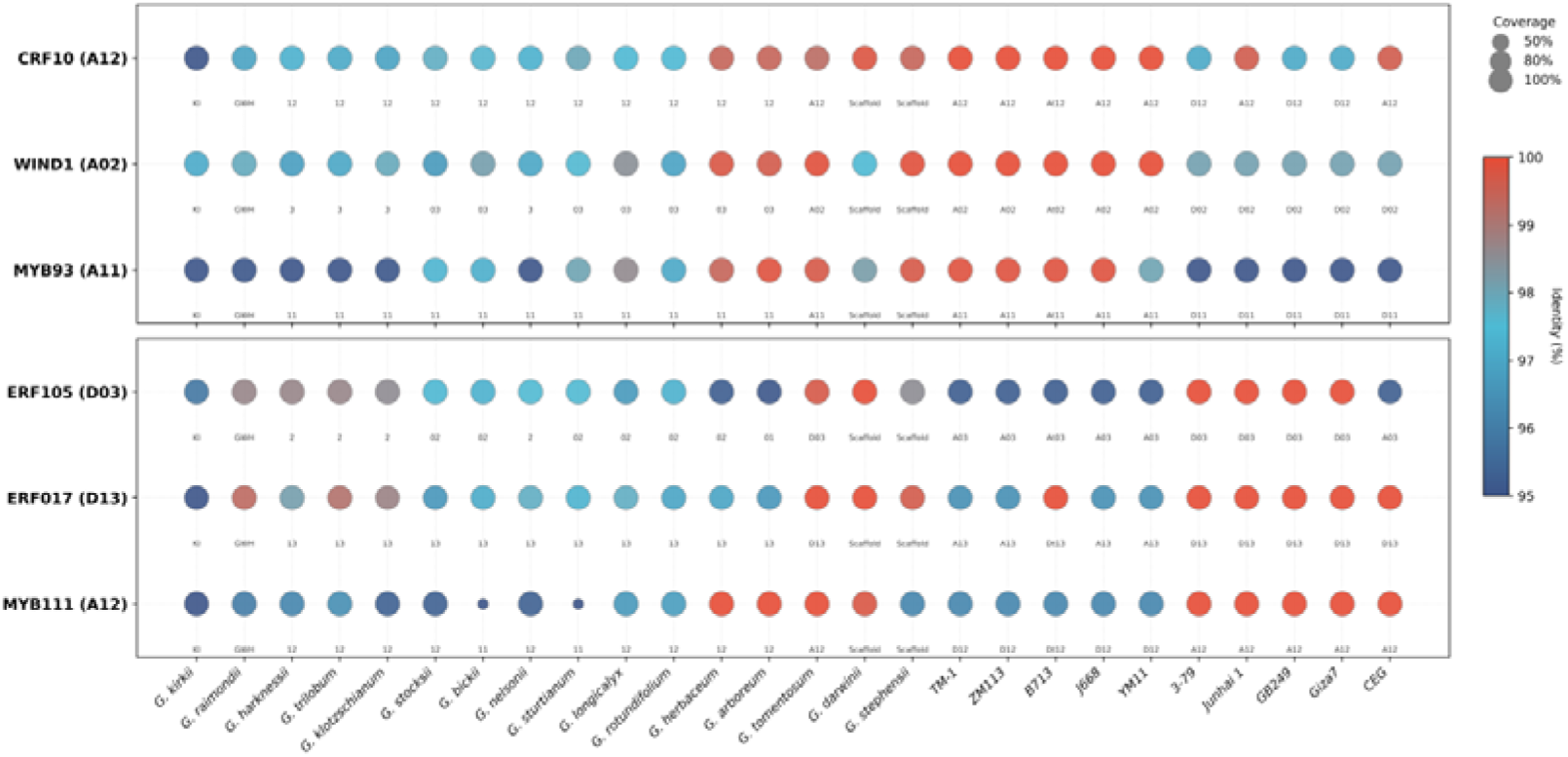
Phylogenomic footprinting of six RSR candidate genes across 26 Gossypium pan-genomes. Bubble plots display the presence and sequence identity of target genes in diploids, wild tetraploids, and cultivars, validating the population-wide universality of the RSR pattern. (a) Upland Cotton Specific Retention Group (Group A): Includes *MYB93* (A11), *WIND1* (A02), and *CRF10* (A12). These genes maintain high sequence identity (red bubbles) and are located on the A-subgenome in all Upland cotton germplasms (e.g., TM-1, J668, ZM113). In contrast, in all Sea Island cotton germplasms (e.g., 3-79, H7124), the A-subgenome copies are lost, and only D-subgenome homeologs with lower sequence identity (blue bubbles) are detected. (b) Sea Island Cotton Specific Retention Group (Group B): Includes *MYB111* (A12), *ERF105* (D03), and *ERF017* (D13). These genes are highly conserved in Sea Island cotton populations (red bubbles) but appear as low coverage or only detectable homeologs in Upland cotton populations. Bubble size represents query Coverage, and color represents sequence Identity. The bottom axis labels chromosomal positions, revealing the drift from “orthologs” to “paralogs/homeologs."

"Alternative Detection” of Homeologs Detects Subgenome-Specific Loss: In the “Upland-Retained Group” (Fig. 6a, e.g., WIN*D1*), all Upland cotton accessions (TM-1, J668, etc.) showed high-identity matches located on chromosome A02 (red bubbles, Identity > 99%), indicating that this gene is core and conserved within the Upland cotton population. However, in all Sea Island cotton accessions (3-79, H7124, Giza7, etc.), BLAST could only detect lower-identity matches located on chromosome D02 (blue bubbles, Identity ∼98%). This “chromosomal drift” from A02 to D02 is crucial—it implies that the original A-subgenome functional copy (*WIND1*) in the Sea Island cotton genome has been completely lost, forcing the probe to “settle for the second best” and match the D-subgenome homeolog. This population-wide loss pattern confirms that the absence of *WIND1* is a species characteristic of Sea Island cotton, rather than a mutation in individual varieties.

Collective Silencing of “Quality Genes” in Upland Population: Conversely, in the “Sea Island-Retained Group” (Fig. 6b, e.g., *MYB111*), the Sea Island cotton population consistently retained the intact gene on chromosome A12. However, in the Upland cotton population, this locus appeared as extremely low coverage (small bubbles) or homeologous drift. This further increases the likelihood that Upland cotton, in pursuit of early maturity and high yield, “purged” this fiber elongation factor at the whole-population level during domestication. In summary, pan-genomic evidence indicates that the RSR mechanism is not accidental genetic drift but a “core genetic imprint” solidified in the genomes of the two species after long-term domestication selection. This highly consistent “retention vs. loss” pattern between species provides a solid theoretical basis for improving Upland cotton using wild germplasm resources or the Sea Island cotton gene pool.

## 4. Discussion

### 4.1 Reciprocal Selective Retention (RSR): A New Genomic Evolutionary Perspective Breaking the “Yield-Quality” Negative Correlation

The negative correlation between “high yield” (typified by Upland cotton) and “high quality” (typified by Sea Island cotton) has long been regarded as an insurmountable bottleneck in cotton breeding, often attributed to linkage drag or physiological energy constraints (Yang et al., 2026). However, our multi-omics integrated analysis offers an alternative genomic explanation: the Reciprocal Selective Retention (RSR) evolutionary model. Unlike the gradual accumulation of Single Nucleotide Polymorphisms (SNPs), RSR reveals a radical “genomic subtraction” strategy, where species actively eliminated genetic modules inconsistent with their survival strategies (Olson, 1999).

Specifically, this model reveals the significantly divergent evolutionary logics adopted by the two species: (1) Upland Cotton’s “High-Yield Robustness Module": To support its superior yield potential, Upland cotton specifically retained *CRF10* (Cytokinin Response Factor, potentially regulating sink capacity and biomass) (Rashotte et al. 2006), *WIND1* (cell regeneration factor) (Iwase et al. 2011), and MYB93 (root system architecture factor) (Gibbs et al. 2014). These genes collectively construct a genetic chassis integrating “strong acquisition (roots), strong sink capacity (CRF), and strong repair (WIND)". (2) Sea Island Cotton’s “Quality Specialization Module": As an evolutionary cost, Sea Island cotton lost the aforementioned broad-adaptability genes, instead specifically retaining *MYB111* (maintaining high flavonoid levels to finely tune/delay the end of elongation, thereby achieving longer fibers) (Stracke et al. 2007; Tan et al. 2013) and *ERF105/ERF017* (stress defense factors) (Bolt et al. 2017). We speculate that these ERF members may act as “environmental buffers” during the prolonged fiber development window, providing necessary physiological homeostasis protection for the precise synthesis of high-quality fibers. This asymmetric “hardware” solidification on homologous chromosomes constitutes the key genomic structural basis for the phenotypic trade-off between the two species.

### 4.2 Transcriptome “Phase Inversion": A Developmental Trade-off of “Time” for “Quality"

Our comparative transcriptomic landscape reveals an intriguing paradox in cotton domestication: vegetative organs are highly conserved, while reproductive organs undergo drastic remodeling. This phenomenon implies that the cotton genome is subject to strong evolutionary constraints to maintain basic life activities (transcriptional chassis) on one hand, while possessing extremely high plasticity in key traits like fiber development on the other.

The most significant discovery is the “Phase Inversion” of the transcriptional program during fiber development. The “delayed elongation” network specifically activated by Sea Island cotton at 20 DPA is essentially a strategy of “trading time for quality.” Through the sustained expression of key factors like *MYB111*, Sea Island cotton delays the initiation of secondary wall thickening, thereby gaining a longer elongation window, which is consistent with previous physiological observations (Avci et al., 2013). In contrast, the transcriptional profile of Upland cotton reflects an “efficiency-first” strategy: the physical loss of *MYB111* leads to the premature termination of the elongation program, forcing cells to rapidly enter the secondary wall filling stage (the blue dominant peak at 25 DPA). Although this “precocious mode” limits fiber length, it significantly shortens the growth cycle and increases boll weight (biomass) (Haigler et al. 2012), perfectly fitting the demand for “high lint percentage and short accumulated temperature” in extensive cultivation.

### 4.3 Evolutionary Constraints and Adaptive Differentiation: Evolutionary Trade-off between “Resource Acquisition” and “Sink Development Timing"

Our comparative transcriptomic landscape and pan-genomic analysis further reveal the co-evolutionary logic of “resource acquisition (Source)” and “sink capacity construction (Sink)” during cotton domestication (White et al., 2016).

First is the “underground cornerstone” for building efficient environmental adaptability. Our data indicate that TM-1 exhibits a significant expansion of specific genes in roots and specifically retains *MYB93*. Previous studies have shown that the homolog of *MYB93* in Arabidopsis acts as a negative regulator of lateral root development (Gibbs et al., 2014). The retention of this “suppressor” by Upland cotton may not be accidental but an optimization strategy for Root System Architecture (RSA): limiting ineffective proliferation of shallow lateral roots to promote deep rooting of the main root or improve water and fertilizer use efficiency (Lynch, 1995). Compared to the sensitivity of Sea Island cotton roots to specific environments, this root remodeling based on genomic specific retention provides a foundation for Upland cotton to establish a robust nutrient absorption network in diverse environments, offering sufficient material and energy support for the subsequent reproductive burst (Hu et al., 2019).

Secondly, supported by a robust “Source", Upland cotton achieved an explosive expansion of “Sink Capacity". The core of the yield difference lies in the increase in boll number per plant and seed number per boll, which mainly depends on the number of locules. Upland cotton typically has 4-5 locules, significantly higher than the 3-locule characteristic of Sea Island cotton (Zhang et al., 2015; Viot and Wendel, 2023). Our analysis precisely captured the transcriptomic imprint determining this trait: the fundamental remodeling of the reproductive organ development program. On one hand, TM-1 showed significant specific gene amplification in anthers, which may enhance pollen viability and fertilization efficiency to meet the high demand of multi-locule and multi-ovule fertilization for male gametes; on the other hand, our common gene analysis revealed that the pistil—the maternal tissue determining carpel fusion and locule number—is the most drastically differentiated floral organ between the two species (common gene proportion < 5%) (Fig. 2). This highly specific transcriptional profile suggests that Upland cotton may have broken the genetic limit of “3 locules” in Sea Island cotton by reconstructing the meristem maintenance pathway or floral organ identity gene network, thereby establishing a high sink capacity architecture of “4-5 locules", which represents a significant advantage for yield.

### 4.4 Implications for Molecular Module Breeding: Reassembling “High-Yield Chassis” and “Quality Modules"

The proposal of the RSR model provides novel insights for cotton precision breeding. Since the separation of “high yield” and “high quality” stems from the physical loss of specific gene modules, future breeding directions should not merely be hybrid screening but “Modular Assembly” based on pan-genomic information.

Strategy 1: Improving the adaptability of Sea Island cotton. Addressing the bottleneck of Sea Island cotton being “difficult to grow", we can use gene editing or introgression breeding to “replenish” the Upland cotton-specific *MYB93-WIND1* root module into Sea Island cotton. Given the key roles of *WIND1* and *MYB93* in cell regeneration and root development (Iwase et al., 2011; Gibbs et al., 2014), this strategy is expected to reshape the underground architecture of Sea Island cotton, enhancing its environmental adaptability and transforming it from a “noble crop” into a more widely promotable variety.

Strategy 2: Breaking the quality ceiling of Upland cotton. Addressing the limitation of fiber length in Upland cotton, we can attempt to introduce the *MYB111* delay module from Sea Island cotton. Previous studies have confirmed that flavonoid metabolism is closely related to fiber elongation (Tan et al., 2013), and *MYB111* is a key regulator of this pathway (Stracke et al., 2007). However, it should be noted that simple replenishment may lead to late maturity. Therefore, using fiber-specific promoters (such as SCW promoters) to finely tune its expression window (Huang et al., 2021) to moderately extend the elongation phase without significantly prolonging the total growth period will be key to creating new “high-yield and high-quality” Upland cotton varieties.

## 5. Conclusion

This study systematically dissected the genetic basis of the “yield-quality” phenotypic trade-off between Upland and Sea Island cotton by integrating multi-tissue transcriptomics and pan-genomic structural variation analysis. We found that: (1) The two species share a conserved transcriptional chassis but achieve functional specialization through asymmetric expansion of gene repertoires in specific tissues (Upland roots/anthers vs. Sea Island fibers). (2) The “Phase Inversion” of the transcriptional program during fiber development is key to fiber quality differences, with Sea Island cotton trading for a longer elongation window via a delayed expression strategy. (3) The “high yield” characteristic of Upland cotton benefits from its specifically retained adaptive genetic modules. For instance, the retained *MYB93* and *WIND1* may have optimized root architecture and environmental adaptability, building a robust “underground resource acquisition” system to support the explosive expansion of the above-ground reproductive sink. (4) This process is driven by the Reciprocal Selective Retention (RSR) mechanism at the genomic level, i.e., the complementary loss of key members of MYB and AP2/ERF families on homologous chromosomes (e.g., Upland cotton discarding *MYB111*, Sea Island cotton discarding *WIND1*). In summary, the RSR mechanism reveals the wisdom of “genomic subtraction” in crop domestication. Future breeding work should focus on the recombination of superior functional modules across species (such as “stress-resistant roots” and “delayed elongation") to break the long-standing bottleneck between yield and quality.

## CRediT authorship contribution statement

**YF Xu**: Conceptualization, Methodology, Software, Formal analysis, Investigation, Data curation, Writing – original draft, Visualization. **Zaoyang Gong**: Validation, Data curation. **Qinglin Shen**:Validation, Investigation. **Rongzheng Zhao**: Data curation, Software. **Chuankang Cheng**:Validation, Investigation. **Wanting Su**: Data curation, Visualization. **Yibing Li**: Validation, Investigation. **Xingrui Yang**: Data curation. **Xin Ruan**: Investigation. **Fengyun Li**: Validation.

**Kai Guo**: Resources, Investigation. **Dajun Liu**: Resources, Investigation. **Xueying Liu**: Supervision, Writing – review & editing. **Zhonghua Teng**: Supervision, Writing – review & editing. **Fang Liu**:Supervision, Writing – review & editing. **Zhengsheng Zhang**: Resources, Supervision. **Yanchao Xu**: Conceptualization, Methodology, Software, Supervision, Writing – review & editing. **Dexin Liu**: Funding acquisition, Project administration, Supervision, Writing – review & editing.

## Declaration of Competing Interest

The authors declare that they have no known competing financial interests or personal relationships that could have appeared to influence the work reported in this paper.

## Acknowledgements

This work was supported by the National Key Research and Development Program of China (Grant No. 2024YFD1200300), the National Natural Science Foundation of China (Grant No. 31701471), and the State Key Laboratory of Cotton Bio-breeding and Integrated Utilization (Grant No. CB2025A13). We also acknowledge the High Performance Computing (HPC) clusters at Southwest University for their support.

## Data availability

Data will be made available on request

## Notes

### Competing Interest Statement

The authors have declared no competing interest.

### Summary of Updates

This revision includes the following significant updates: New Analysis Integrated: We have added a comprehensive genome-wide analysis of the AP2/ERF transcription factor family to complement the MYB analysis. This expands the core findings to a "3-vs-3" reciprocal selective retention pattern (involving MYB111, ERF105, ERF017 vs. MYB93, WIND1, CRF10), providing stronger evidence for the RSR model. Figures Reorganized: Figures have been updated and merged to better illustrate the transcriptomic phase inversion and genomic structural variations. Logic Refined: The manuscript text has been revised to clarify the logical flow and the evolutionary mechanisms driving the yield-quality trade-off. Title Updated: The title has been updated to "Transcriptomic Phase Inversion and Reciprocal Selective Retention Drive the Yield-Quality Trade-off in Cultivated Cotton" to better reflect the broadened scope.

