## Supplementary Materials for "Multi-tissue Transcriptomic and Pan-genomic Analyses Reveal Reciprocal Selective Retention Driving the Phenotypic Trade-off in Gossypium hirsutumand Gossypium barbadense"

**Supplementary Table S1. Summary of genomic information for the 26 Gossypium accessions used in pan-genomic analysis.**

| Species | Number of Genes | Genomic Group | Genome Size | Reference |
| --- | --- | --- | --- | --- |
| G.herbaceum | 41642 | A1 | 1.43G | Wang et al. (2022) |
| G.arboreum | 41778 | A2 | 1.50G | Wang et al. (2021) |
| G.sturtianum | 41308 | C1 | 1.77G | Wang et al. (2022) |
| G.harknessii | 36314 | D2-2 | 692MB | Xu et al. (2024) |
| G.klotzschianum | 43254 | D3-k | 771MB | Xu et al. (2024) |
| G.raimondii | 53167 | D5 | 747MB | Huang et al. (2024) |
| G.gossypioides | 39601 | D6 | 704MB | Xu et al. (2024) |
| G.trilobum | 37605 | D8 | 693MB | Xu et al. (2024) |
| G.stocksii | 40321 | E1 | 1.34G | Wang et al. (2022) |
| G.longicalyx | 41633 | F1 | 1.11G | Wang et al. (2022) |
| G.bickii | 40857 | G1 | 1.49G | Wang et al. (2022) |
| G.rotundifolium | 41590 | K2 | 2.27G | Wang et al. (2021) |
| G.kirkii | 36680 | Outgroup | 520MB | Grover et al. (2019) |
| TM-1 | 79642 | AD1 | 2.14G | Hu et al. (2025a) |
| B713 | 78475 | AD1 | 2.17G | Perkin et al. (2021) |
| J668 | 74637 | AD1 | 2.24G | Xu et al. (2025) |
| YM11 | 84720 | AD1 | 2.18G | Wang et al. (2023) |
| ZM113 | 78467 | AD1 | 2.18G | Hu et al. (2025b) |
| 3-79 | 76754 | AD2 | 2.13G | Chang et al. (2024) |
| Junhai1 | 75774 | AD2 | 2.11G | Meng et al. (2025) |
| Giza7 | 76012 | AD2 | 2.08G | Meng et al. (2025) |
| GB249 | 71549 | AD2 | 2.09G | Meng et al. (2025) |
| CEG | 75755 | AD2 | 2.10G | Meng et al. (2025) |
| G.tomentosum | 78281 | AD3 | 2.06G | Chen et al. (2020) |
| G.darwinii | 78303 | AD5 | 2.06G | Chen et al. (2020) |
| G.stephensii | 74970 | AD7 | 2.15G | Peng et al. (2022) |
| TM-1 | 79642 | AD1 | 2.14G | Hu et al. (2025a) |
| B713 | 78475 | AD1 | 2.17G | Perkin et al. (2021) |
| J668 | 74637 | AD1 | 2.24G | Xu et al. (2025) |

Note: The table lists the genomic statistics for the 26 accessions used in the phylogenomic and microsynteny analyses. Species: TM-1 (G. hirsutum) and 3-79 (G. barbadense) serve as the primary reference genomes for Upland and Sea Island cotton, respectively. Genomic Group: Classifications include diploid ancestors (A, C, D, E, F, G, K genomes), the outgroup (G. kirkii), and allotetraploids (AD1–AD7). Genome Size: Sizes are indicated in Gigabases (Gb) or Megabases (Mb). Reference: Citations correspond to the original genome assembly publications; full bibliographic details are provided in the References section of the main text. Data were retrieved from CottonGen ([https://www.cottongen.org](https://www.cottongen.org" \t "_blank)) and NCBI.

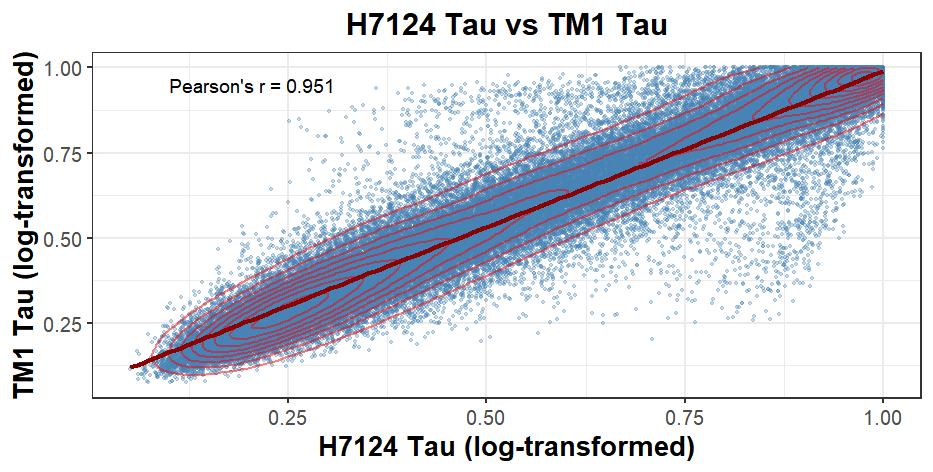

**Supplementary Fig. 1. Evolutionary conservation of tissue specificity between Upland cotton (*G. hirsutu*m) and Sea Island cotton (*G. barbadense*).** A scatter plot comparing the Tissue Specificity Index (Tau) of orthologous gene pairs between H7124 (x-axis) and TM-1 (y-axis). The red line represents the linear regression fit. The high Pearson correlation coefficient (*R* = 0.951) indicates a high degree of conservation in the expression breadth of most genes. Points deviating from the diagonal (outliers) represent genes with divergent tissue specificity between the two species.

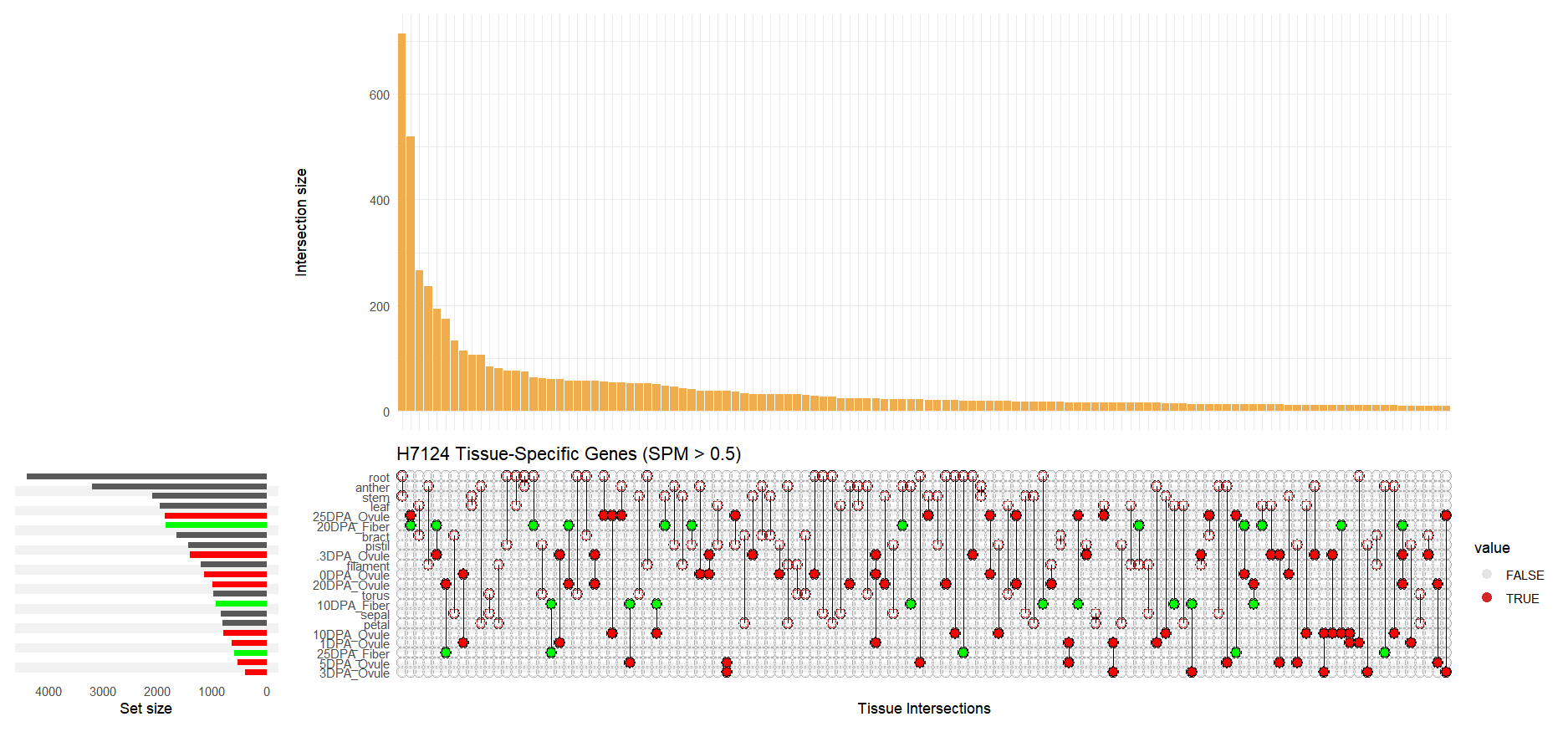

**Supplementary Fig. 2. Intersection and distribution landscape of tissue-specific genes in Sea Island cotton (H7124) (UpSet Plot).** This plot displays the number and intersections of specifically expressed genes (SPM > 0.5) across tissues in H7124. Vertical bars represent the size of specific intersections (here focusing primarily on single-tissue specific genes). Note that in the late stages of fiber development (especially 20 DPA and 25 DPA Fiber) and ovule maturation (25 DPA Ovule), H7124 still maintains a vast repertoire of specific genes, forming a sharp contrast with Upland cotton.

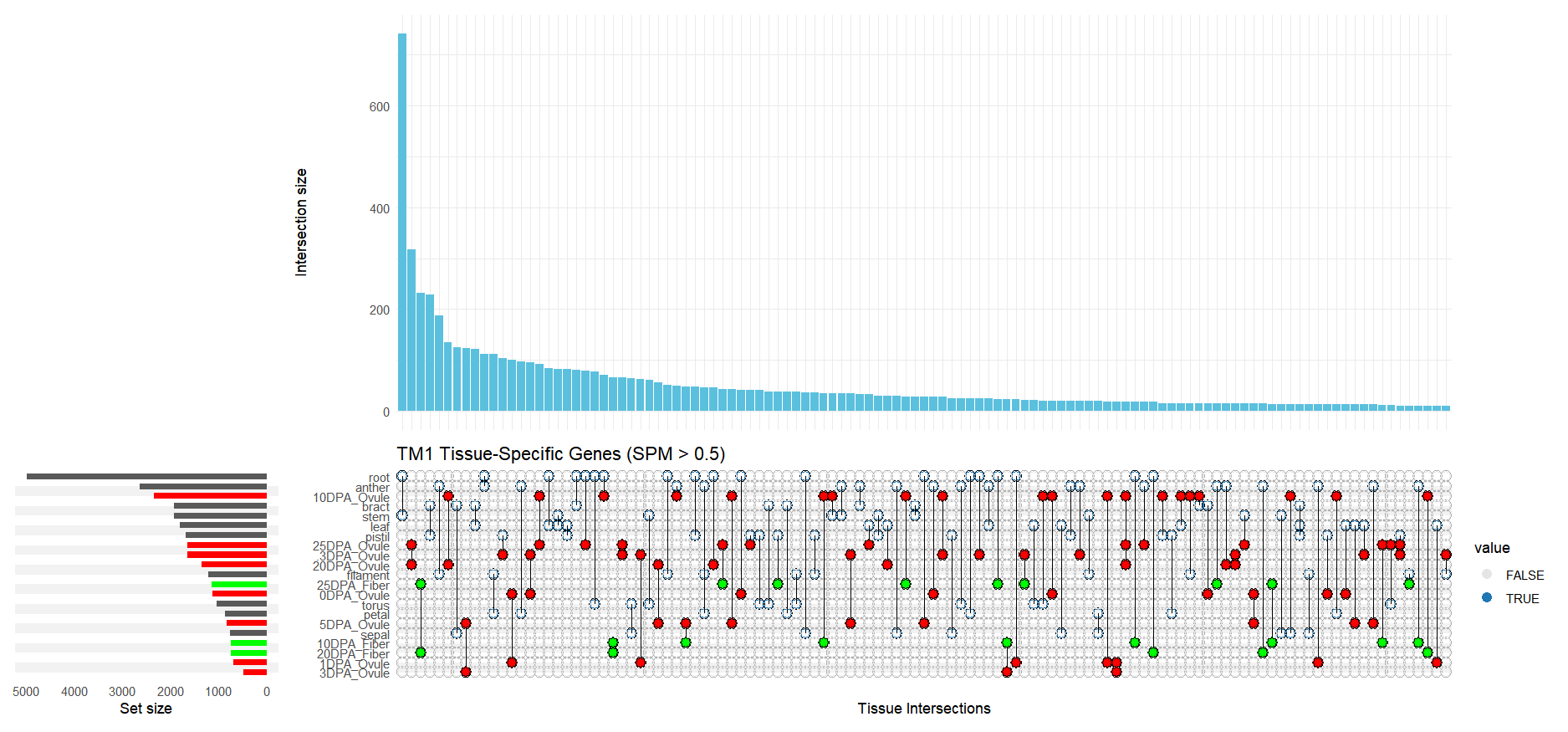

**Supplementary Fig. 3. Intersection and distribution landscape of tissue-specific genes in Upland cotton (TM-1) (UpSet Plot).** This plot shows the distribution of specifically expressed genes across tissues in TM-1. Unlike Sea Island cotton, the most distinct feature of TM-1 is the explosive expansion of root-specific genes, constituting the tallest bar in the plot. Conversely, during the middle to late stages of fiber development (10–25 DPA), the number of specific genes shows a marked contraction.

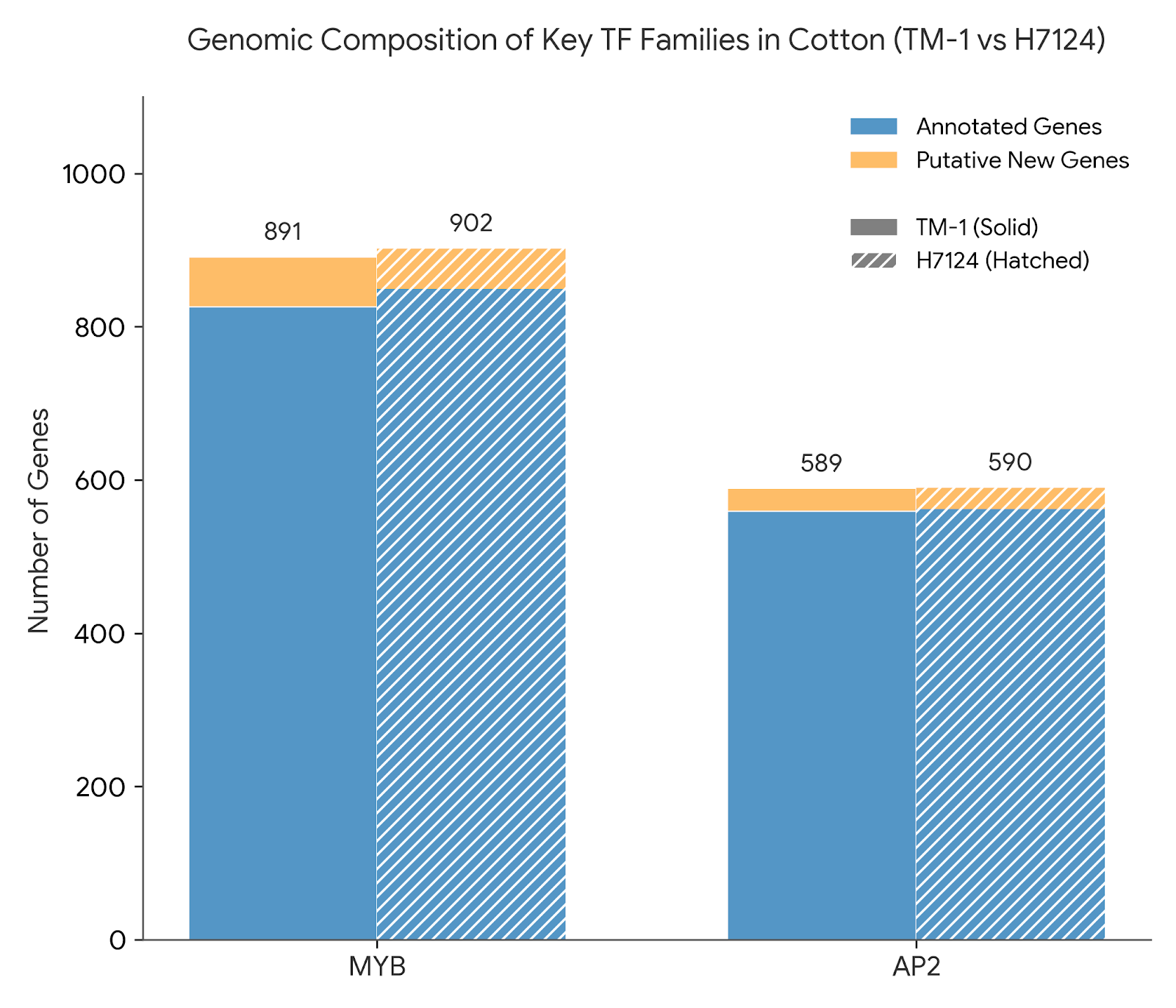

**Supplementary Fig. 4. Genome-wide composition statistics of key transcription factor families (MYB and AP2) in Upland cotton (TM-1) and Sea Island cotton (H7124).** Stacked bar charts display the total number and classification of MYB and AP2 gene family members in both species. Blue bars (Annotated Genes) represent genes originally annotated in the reference genomes, while orange bars (Putative New Genes) represent loci newly identified via the Bitacora pipeline. Despite the vast divergence in agronomic traits between the two species, family sizes at the whole-genome level exhibit high evolutionary conservation (e.g., MYB: 891 vs 902; AP2: 589 vs 590). This stability in macroscopic quantity strongly implies that phenotypic differentiation is driven not by drastic expansion or contraction of gene family sizes (Dosage Effect), but more likely by functional replacement or structural variation of specific key members.

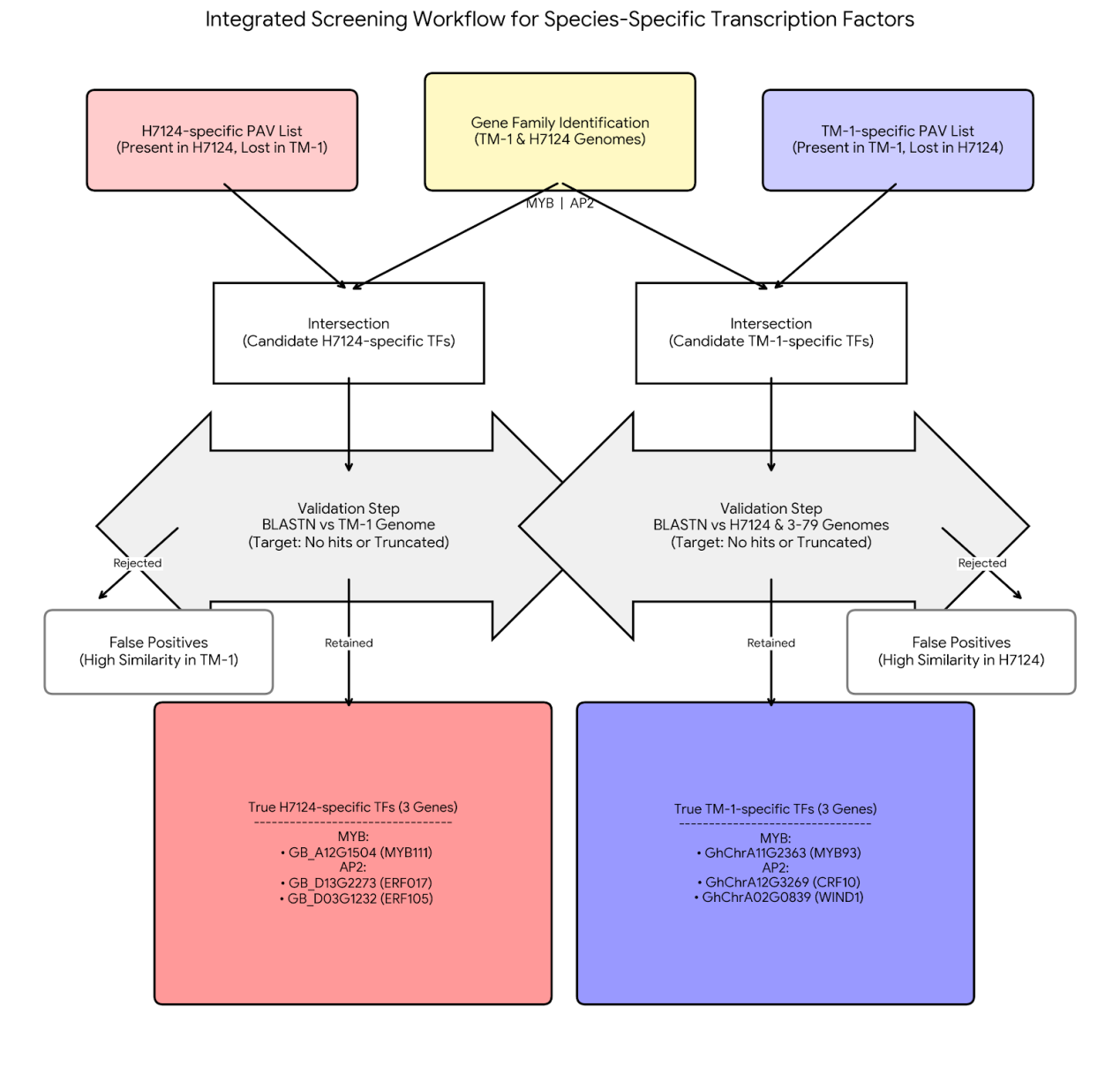

**Supplementary Fig. 5. "PAV Intersection-Reciprocal Validation" screening workflow for identifying high-confidence species-specific transcription factors.** The schematic illustrates the comprehensive strategy for pinning down six key candidate genes at the whole-genome level. (1) Initial Screening: The whole-genome Presence/Absence Variation (PAV) list was intersected with identified MYB and AP2 family members to obtain a candidate list. (2) Reciprocal Validation: Strict BLASTN reciprocal alignment was implemented. Only genes with no homologous fragments detected or exhibiting severe sequence truncation in the counterpart genome were retained to thoroughly eliminate false positives caused by missing annotations. (3) Final Output: Three H7124-specific retained genes (red boxes) and three TM-1-specific retained genes (blue boxes) were successfully identified, constituting the core targets for subsequent structural variation analysis.

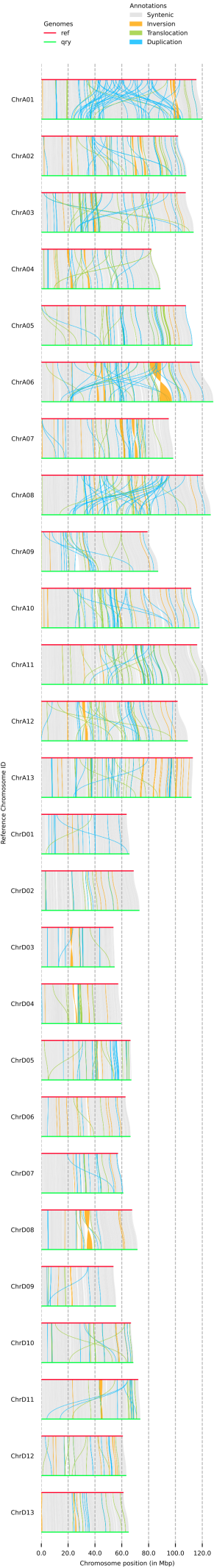

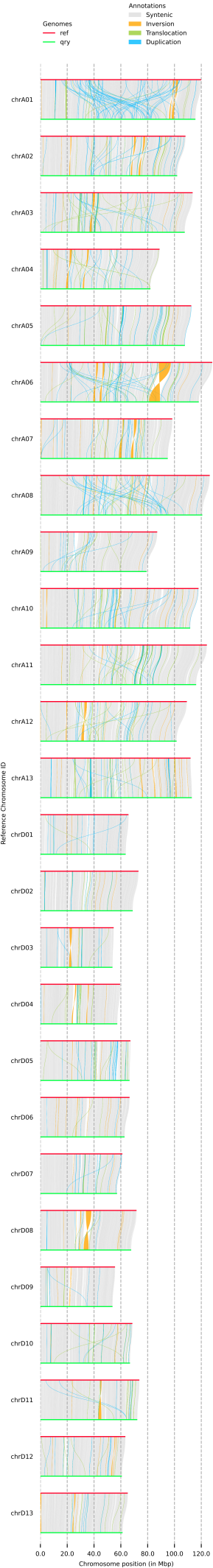

**Supplementary Fig. 6. Whole-genome structural variation map referenced to Sea Island cotton (Ref: H7124 vs Query: TM-1).** Whole-genome structural variation landscape of all chromosomes (A01-A13 and D01-D13) identified by SyRI and visualized by plotsr. (a) The upper chromosome backbone represents the reference genome H7124, and the lower represents the query genome TM-1. (b)Gray bands show extensive synteny between the two species. Colored blocks mark structural variations (e.g., inversions, translocations) in TM-1 relative to H7124. (c) This perspective highlights the authenticity of H7124-specific retained genes (Group 2: MYB111, ERF105, ERF017). On the H7124 coordinate axis, these gene loci are structurally intact, whereas the corresponding TM-1 regions display physical deletion (NOTAL) or high sequence divergence (HDR).

**Supplementary Fig. 7.** Whole-genome structural variation map referenced to Upland cotton (Ref: TM-1 vs Query: H7124). Results of the reciprocal SyRI whole-genome structural variation analysis. (a) The lower chromosome backbone represents the reference genome TM-1, and the upper represents the query genome H7124. (b) This perspective anchors the genomic positions of TM-1 specific retained genes (Group 1: MYB93, WIND1, CRF10). In the TM-1 reference coordinates, these genes are located within highly conserved syntenic blocks, while the corresponding H7124 regions are identified as large-fragment deletions or sequence collapse. This reciprocal validation strategy provides chromosomal-level evidence for the loss of these genes.
